# Strain-specific changes in nucleus accumbens transcriptome and motivation for food reward in mice exposed to maternal separation

**DOI:** 10.1101/2023.04.06.535855

**Authors:** Simon Benoit, Mathilde Henry, Sara Fneich, Alexia Mathou, Lin Xia, Aline Foury, Mélanie Jouin, Claudine Junien, Lucile Capuron, Luc Jouneau, Marie-Pierre Moisan, Cyrille Delpierre, Anne Gabory, Muriel Darnaudery

## Abstract

Adversity in childhood exerts enduring effects on brain and increases the vulnerability to psychiatric diseases. It also leads to a higher risk for eating disorders and obesity. We hypothesised that neonatal stress in mice affects motivation to obtain palatable food in adulthood and changes gene expression in reward system. Male and female pups from C57Bl/6J and C3H/HeN mice strains were subjected to a daily maternal separation (MS) protocol from PND2 to PND14. In adulthood, their motivation for palatable food reward was assessed in operant cages. Compared to control mice, male and female C3H/Hen mice exposed to MS significantly did more lever presses to obtain palatable food especially when the effort required to obtain the reward is high. Transcriptional analysis reveals 375 genes differentially expressed in the nucleus accumbens of male MS C3H/HeN mice compared to the control group, some of these being associated with the regulation of the reward system (e.g. *Gnas, Pnoc*). Interestingly, C57Bl/6J mice exposed to MS did not show any alteration in their motivation to obtain a palatable reward nor significant changes in gene expression in the nucleus accumbens. In conclusion, neonatal stress produces lasting changes in motivation for palatable food in C3H/HeN offspring but has no impact in C57Bl/6J offspring. These behavioural alterations are accompanied by drastic changes in gene expression specifically within the nucleus accumbens, a key structure in the regulation of motivational processes.

## 1 Introduction

In human, early-life adversity during infancy influences the development of child and exerts long-lasting effects on physiological functions and vulnerability to psychiatric disorders, notably depression, anxiety disorders, and substance abuse (Heim et al., 2008; Hughes et al., 2017; Teicher et al., 2022). Early-life adversity produces numerous physiological abnormalities including hypothalamic-pituitary-adrenocortical (HPA) axis hyperactivity, low grade inflammation, and affects brain areas involved in the regulation of cognitive and emotional processes such as the medial prefrontal cortex (mPFC), amygdala, hippocampus, and ventral striatum (Nusslock and Miller, 2016; Danese and J Lewis, 2017). Additionally, early-life adversity exacerbates vulnerability to obesity in adult subjects (Danese and Tan, 2014), and this effect could be at least partially attributed to poor feeding habits and/or exacerbated motivation for high caloric density food (Mason et al., 2013; Osadchiy et al., 2019). However, despite an important literature on the impact of early-life adversity on neuropsychiatric vulnerability, its impact on food motivation remains less explored (Miller and Lumeng, 2018).

In rodents, chronic maternal separation (MS) has been used as a *proxy* of early-life adversity. MS effects on emotional behaviours have been extensively documented in rats (Franklin et al., 2012; Rincel and Darnaudéry, 2020). These effects include cognitive impairments, exacerbated anxiety- and depressive-like behaviour as well as anhedonia associated with HPA axis alterations (Sánchez et al., 2001; Schmidt et al., 2011; Rincel et al., 2016). Perinatal stress, both during prenatal and postnatal period, is also associated with metabolic disturbances (Vallée et al., 1996; Lesage et al., 2004; Ilchmann-Diounou et al., 2019) and a higher sensitivity to diet-induced obesity (Romaní-Pérez et al., 2017). Finally, a growing body of evidence suggests that early-life stress impairs reward processes (Novick et al., 2018). While MS has been associated with reduced sucrose preference suggestive of an anhedonia phenotype (Rincel and Darnaudéry, 2020; Birnie et al., 2023), early-life adversity was found to exacerbate drug intake in rodents as well as in humans (Nylander and Roman, 2013; Walters and Kosten, 2019). MS has been associated with disruptions of the reward dopaminergic system (Rodrigues et al., 2011; Romaní-Pérez et al., 2017; Wendel et al., 2021). Recent studies showed long lasting transcriptomic changes in the ventral tegmental area and in the nucleus accumbens (NAc) of mice exposed to MS combined with unpredictable chronic mild stress or social defeat in adulthood (Peña et al., 2017, 2019). However, the impact of MS on mice motivation for palatable food is still unknown. This is an important issue given that reward circuit alterations have been recurrently reported in the literature in both animal models and humans exposed to early-life adversity (Wendel et al., 2021).

The aim of the present study was then to determine the impact of MS on motivation for palatable food in female and male mice offspring. Since mice are particularly resilient to early-life stress procedures (Tractenberg et al., 2016; Tan et al., 2017), we studied the effects of MS in C57Bl/6J and C3H/HeN mice, two mouse strains used for MS paradigms (Franklin et al., 2010; Peña et al., 2017; Riba et al., 2018; Rincel et al., 2019). To further characterise the brain changes associated with MS, we also examined gene expression in the hypothalamus, the NAc, and the mPFC.

## 2 Materials and methods

### 2.1 Animals

All experiments were carried out in accordance with French (Directive 87/148, Ministère de l’Agriculture et de la Pêche) and European (Directive 2010/63/EU, 2010 September 22th) legislation and approved by Institutional Regional Committee for animal experimentation (agreement #5012050-A). C57Bl/6J and C3H/HeN mice were obtained from Janvier Labs (Le Genest, Saint-Isle, France) and housed under standard laboratory conditions (23 ± 1°C; 12 h/12 h light/dark cycle; lights on at 7 a.m.; food and water *ad libitum*). After one week of habituation, two nulliparous female mice (11 weeks old) were placed with one male from the same strain during one week for breeding and then pregnant dams were single-housed in polycarbonate cages (48 × 26 × 21 cm) throughout gestation and lactation. The day of delivery was designated postnatal day 0 (PND0). At PND1, pups from all litters were pulled, sexed, and weighed. Two litters with abnormal number of pups (< 3) or sex-ratio (only females) were excluded from the study. Litters were assigned to Maternal Separation (MS, C57Bl/6J, n = 7; C3H/HeN, n = 7) or control (C57Bl/6J, n = 6; C3H/HeN, n = 6) groups.

### 2.2 Maternal separation combined with chronic unpredictable maternal stress

MS was carried out from PND2 to PND14 (180 min daily) (Rincel et al., 2016, 2019) and started randomly at 8:30, 9:00, 9:30, 10:15, 10:30 or 11:00 to minimize habituation. During separation sessions, pups were individually separated and kept at 32 °C ± 2. During the separation, dams were exposed to a chronic unpredictable stress protocol (PND2: no bedding; PND3: sodden bedding; PND4: tilted cage 45°; PND5: soiled rat bedding; PND6: sodden bedding; PND7: no bedding; PND8: tilted cage 45°; PND9: no bedding; PND10: tilted cage 45°; PND11: sodden bedding; PND12: no bedding; PND13: soiled rat bedding; PND14: forced swim test (adapted from (Franklin et al., 2010; Rincel et al., 2019)). Control pups were left undisturbed with the dams. All pups were weaned on PND21 and grouped in 4-6 mice per cage by same strain, same sex, and same MS condition.

### 2.3 Behavioural assessment in operant chambers

At least two weeks before the beginning of the behavioural test, the light/dark cycle was inverted (lights off at 7 a.m.) in order to study mice behaviour during the active phase of their cycle. Experiments were conducted during the dark phase between 09:00 and 17:00 h. Male and female offspring’s food motivation for a palatable reward (10% condensed milk in water, 3.25 kcal/g), was assessed at 4-5 months of age in daily 60-min sessions (5 sessions per week) in operant chambers (Imétronic, Pessac, France) equipped with two levers, as previously described (Gueye et al., 2018; Ducrocq et al., 2020). For the habituation and the initial training on fixed-ratio 1, mice were food restricted to 85% of their body weight. Then, animals were fed *ad libitum* throughout the experiment except for the concurrent choice test. *Ad libitum* access to food schedule was used in order to examine motivation for palatable food reward independently of the homeostatic state of the animals. Male and female cohorts were tested separately.

#### 2.3.1 Habituation to the apparatus

Mice were placed into the operant chambers without lever for 30 min and milk reward delivered in the drinking cup every 60 s interval. A dose of milk was distributed only when the previous one had been consumed.

#### 2.3.2 Fixed-ratio 1

Mice were initially trained to press one of the two levers (the active lever) on a fixed-ratio 1 (FR-1) schedule (every active lever press resulted in fluid delivery) to obtain 15 μl of sweetened milk solution in 60-min daily sessions. The delivery of milk was paired with a 4 s cue light above the lever. Responses on the other lever (the inactive lever) were recorded as a measure of non-specific activity. Mice received four FR-1 sessions under mild food deprivation, then mice were tested for an additional FR-1 session in *ad libitum* conditions.

#### 2.3.3 Random-ratio

After FR-1, mice were submitted to random-ratio schedules (RR) in 60-min daily sessions: RR5 (12 sessions), RR10 (3 sessions), and RR20 (12 sessions), with respectively a probability of 1/5, 1/10 or 1/20 to be reinforced after one press on the active lever. RR schedules lead to reinforcement following an unpredictable average number of responses per rewards and result in high and consistent response rates that exceed those obtained with FR or interval schedules.

#### 2.3.4 Progressive-ratio

Following training under RR20 schedule, mice were under a progressive ratio (PR) schedule for 2 daily sessions to assess the motivation for the palatable reward. During PR, the number of lever presses required to earn the next reward increased progressively and is multiplied by 2 (2, 4, 8, 16, 32 etc.) For each reinforced response, the animal received sweetened milk (15 μl). The breakpoint was defined as the last ratio completed.

#### 2.3.5 Devaluation extinction test

Devaluation test allows assessing alteration of the representation of the outcome value. In this test, mice were pre-fed *ad libitum* to either milk or soy-bean oil emulsion Intralipid for 30 min in their home-cage. Immediately after, they were placed into the operant chamber to conduct a lever press test in a 5-min extinction session. Although both levers were introduced, no reward was distributed. Then, mice received 2 days retraining using a RR20 procedure with free access before the second test. This second session was conducted as previously described, however this time, mice were pre-fed with the alternative outcome. The distribution order of the reward is randomly alternated for each mouse from one session to another.

#### 2.3.6 Concurrent choice

Following one session under RR20 schedule with a food deprivation, a concurrent choice 60-min session test was conducted. In this task, fasted mice could press lever for sweet milk, but their standard lab chow was also available on the floor of chamber opposite to the location of the lever. The total number of lever presses and the amount of lab chow consumed were recorded.

### 2.4 Plasma corticosterone

After the completion of the behavioural assessment, two blood samples were collected, one at the beginning of the active phase and one at the beginning of the inactive phase to assess circadian variation of plasma corticosterone levels. Blood was collected by tail nick using EDTA-coated tubes, were centrifuged (4000 rpm, 4°C) for 10 min and stored at -20°C until use. Plasma corticosterone levels were determined with an in-house radioimmunoassay using a highly specific antibody as previously described (Rincel et al., 2016). Cross reactivity with related compound such as cortisol was less than 3%. Intra- and inter-assay variations were less than 10% and less than 15%, respectively.

### 2.5 Metabolic hormones assay

Plasma resistin, leptin and insulin levels were measured in 12 h-fasted mice using a Mouse Metabolic Hormone Magnetic Bead-based immunoassay kits (Millipore Corp, Billerica, MA) according to the manufacturer instructions on samples obtained at the sacrifice.

### 2.6 Culling and samples collection

Mice were 12 h-food deprived and then were deeply anesthetized with isoflurane and culled by decapitation. Blood was collected for metabolic hormones assessment. Whole brains were collected, medial prefrontal cortex (mPFC) and hypothalamus (HT) were dissected; whereas nucleus accumbens (NAc) was punched on frozen slices and store at -80°C until use.

### 2.7 Microarrays

Total mRNA was extracted from mPFC, NAc, and hypothalamus using a TRIzol extraction kit (Invitrogen) according to the manufacturer’s instructions. RNA concentration, purity and integrity were determined using a ND-1000 spectrophotometer (Nanodrop Technologies, Wilmington, DE, USA) and a bioanalyzer (Agilent, Les Ulis, France) (Rincel et al., 2019). Gene expression profiles were performed at the GeT facility (https://www.genotoul.fr/) using Agilent Sureprint G3 Mouse microarrays (8×60K, design 074809) following the manufacturer’s instructions. For each sample, Cyanine-3 (Cy3) labelled cRNA was prepared from 200 ng (mPFC/HT) or 40 ng (NAc) of total RNA using the One-Color Quick Amp Labelling kit (Agilent Technologies, Santa Clara, CA) according to the manufacturer’s instructions, followed by Agencourt RNAClean XP purification (Agencourt Bioscience Corporation, Beverly, Massachusetts). Dye incorporation and cRNA yield were checked using Dropsense™ 96 UV/VIS droplet reader (Trinean, Belgium). Six hundred ng of Cy3-labelled cRNA were hybridized on the Agilent SurePrint G3 Mouse GE microarray slides following the manufacturer’s instructions. Immediately after washing, the slides were scanned on Agilent G2505C Microarray Scanner using Agilent Scan Control A.8.5.1 software and fluorescence signal extracted using Agilent Feature Extraction software v10.10.1.1 with default parameters.

Microarray data and experimental details are available in NCBI’s Gene Expression Omnibus (Edgar et al., 2002) and are accessible through GEO Series accession number GSE222781. Microarray data were analysed using R (R Core Team, 2022) and Bioconductor packages (www.bioconductor.org, v 3.0, (Gentleman et al., 2004) as described in GEO accession

### 2.8 TaqMan low-density arrays

Gene expression was quantified using custom TaqMan low-density arrays (TLDAs, Applied Biosystems). All samples were treated with DNase. The experiment was performed on the @BRIDGE platform (INRAE, Jouy-en-Josas, France) according to the manufacturer’s instructions. Four samples were run on each TLDA card in simplicate. Each sample reservoir on the card was loaded with 100 μl of the reaction mix: cDNA template (600 ng) mixed with TaqMan Gene Expression Master Mix (Applied Biosystems). After centrifugation (twice 1 min at 1200 rpm, Heraeus Multifuge 3S Centrifuge), the wells were sealed with a TLDA Sealer (Applied Biosystems). PCR amplification was performed on the 7900HT Real-Time PCR System (Applied Biosystems) using SDS 2.4 software with standard conditions: 2 min 50 °C, 10 min 94.5 °C, 30 s 97 °C (40 cycles), 1 min 59.7 °C. Threshold cycle (Ct) values were calculated with the ExpressionSuite v1.0.3 software (Applied Biosystems). The detection threshold was set manually for all genes and was the same for each assay in all tissues. Ct = 39 was used as the cut-off above which, expression level was set to 0. On the TLDA array, 96 genes were studied for each sample: 87 target genes and 9 reference genes (Supplementary files F1). Target genes were chosen among the differentially expressed genes found in the microarray experiment in different categories of pathways and biological function. Six on the 9 reference genes were defined as the best reference, using the GeNorm software (Vandesompele et al., 2002). For each sample, Ct[ref] was the mean of the three Ct values of the reference genes. Then, expression level of target genes was calculated as 2 − (Ct[target gene] − Ct[ref]), as previously described (Panchenko et al., 2016).

### 2.9 Statistics

Values are expressed as means ± SEM. All data were analysed using GraphPad version 7.0 software (La Jolla, CA, USA). Student’s t-tests were used to compare body weight, plasma hormone levels, progressive ratio result and chow consumption between control and MS groups. For food motivation in operant chamber, two-way ANOVAs followed by post-hoc analysis to test specific comparisons. were used. P-values < 0.05 were considered statistically significant. Microarray data were analysed using R (R Core Team, 2022) and Bioconductor packages (www.bioconductor.org, v 3.0, (Gentleman et al., 2004). Hierarchical clustering was performed using Pearson’s correlation coefficient as distance function and Ward as linkage method. Partial least squares-discriminant analysis (PLS-DA) was performed for each tissue, with group (Ctrl or stress) as a Y response, using *ropls* package (Thévenot et al., 2015). Transformed signals were mean-centered and divided by the standard deviation of each variable. A model was fitted using the *limma lmFit* function (Smyth, 2005). Pair-wise comparisons of biological conditions were applied using specific contrasts. Probes with Benjamini–Hochberg (BH) false discovery rate (FDR) < 0.05 were considered to be differentially expressed between conditions. Volcano plots were constructed with the *ggplot* function of the R *ggplot2* package. The differentially expressed gene datasets were uploaded into Ingenuity Pathway Analysis software (Qiagen IPA, content version 28820210) and a core analysis was performed, with the Agilent SurePrint G3 Mouse GE microarray as background. The canonical pathways with BH p-values < 0.05 only and the upstream regulators with no flag “bias”, activation z-score < -2 or > 2 and p-value of overlap < 0.05 only were considered. Another analysis was realized using ConsensusPathDB-mouse (Kamburov et al., 2011), with the differentially expressed gene list vs the mus musculus database. KEGG pathways with a q-value < 0.05 were considered. For TLDA, analysis was performed with the R statistical software (R Core Team, 2022). For each gene, we compared the expression values within each maternal group in males and females, in C3H/HeN and C57Bl/6J apart. Pair-wise comparisons of maternal groups were conducted using a permutation test, as implemented in the oneway_test function of the *coin* package in R. For each set of tests (*i*.*e*., all tested genes for a given pair of maternal groups), *p*-values were BH adjusted for multiple testing. Differences were considered significant when *p*-adj < 0.05. The TLDA data clustering was performed using the ClustVis web tool for visualizing clustering of multivariate data (https://biit.cs.ut.ee/clustvis/) (Metsalu and Vilo, 2015).

## 3 Results

### 3.1 Chronic maternal separation impairs C3H/HeN mice’s body weight

The MS paradigm has been used as a *proxy* to model early-life stress in rodents. At the end of the MS procedure, stressed C3H/HeN dams displayed a significant lower body weight compared to the undisturbed C3H/HeN dams (Figure 1B, t_(15)_ = 4.538, p < 0.001), whereas MS only had a minimal impact on C57Bl/6J dams’ weight (Figure 1A, t_(15)_ = 1.937, p = 0.0719). In offspring, both males (Figure 1D, t_(14)_ = 2.518, p < 0.05) and females (Figure S1B, t_(14)_ = 3.986, p < 0.01) C3H/HeN pups and female C57Bl/6J pups (Figure S1A, t_(13)_ = 2.500, p = 0.0266) exhibited a significant decreased body weight at the end of the MS procedure. In contrast, males C57Bl/6J pups were unaffected by MS (Figure 1C). The effect of MS on body weight was maintained until adulthood in C3H/HeN male offspring (Figure 1F, t_(21)_ = 4.035, p < 0.001).

**Figure 1.**
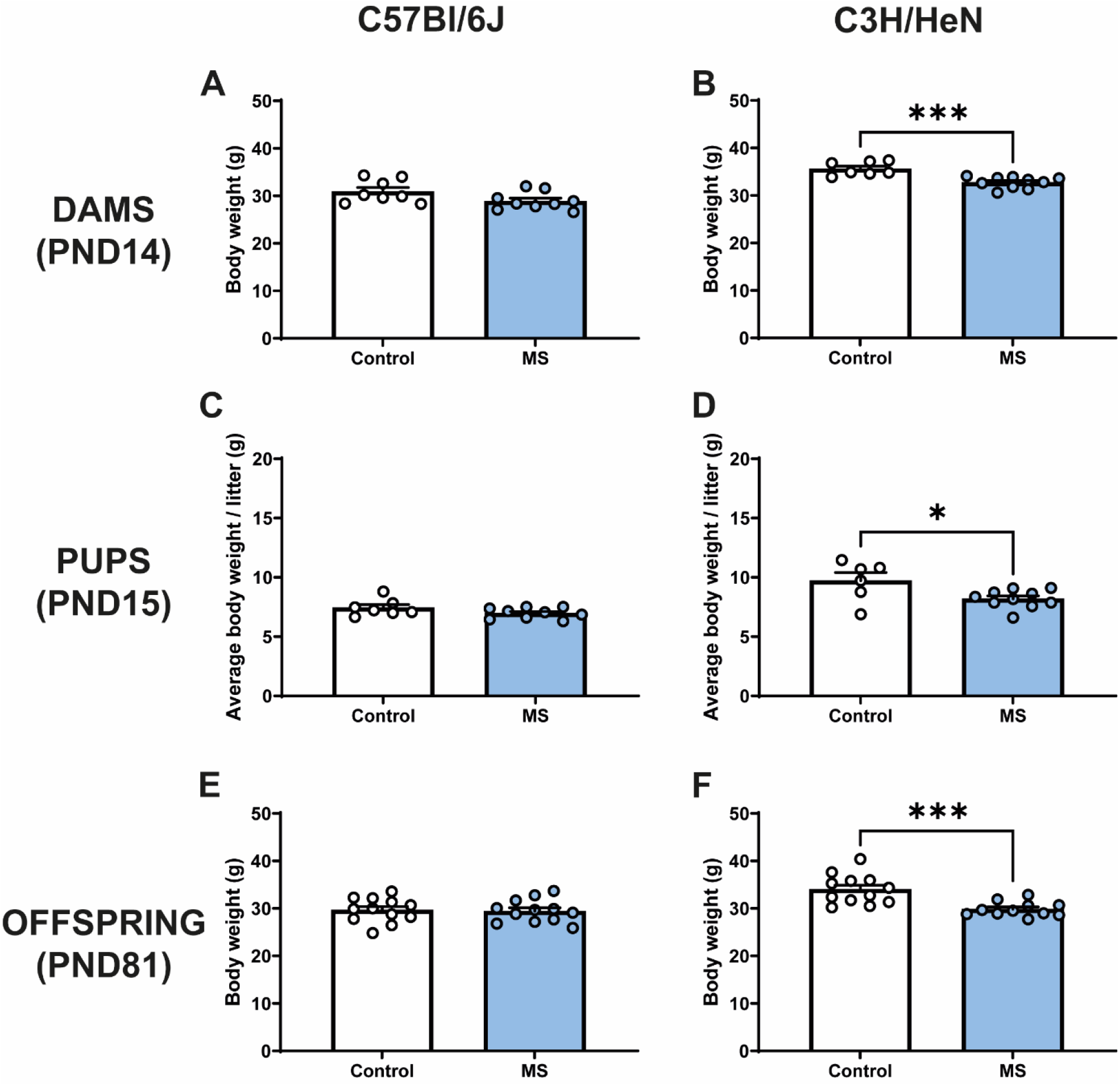
Maternal separation decreases body weight of C3H/HeN in both dams and male offspring. Body weight in Control and MS (A) C57Bl/6J dams at PND14 (Control n = 8, MS n = 9), (B) C3H/HeN dams at PND14 (Control n = 7, MS n = 10), (C) C57Bl/6J male pups at PND15 (Control n = 7, MS n = 9), (D) C3H/HeN male pups at PND15 (Control n = 6, MS n = 10), (E) C57Bl/6J male offspring at PND81 (Control n = 12, MS = 12), and (F) C3H/HeN male offspring at PND81 (Control n = 12, MS n = 12). Data are presented as means ± SEM, *p < 0.05, ****p < 0.001.

### 3.2 Maternal separation increases plasma corticosterone levels in both C57Bl/6J and C3H/HeN strains

The impact of MS on corticosterone and metabolic hormone plasma levels was studied in adult mice. Plasma corticosterone levels were determined at the beginning of dark and light phases. Both C57Bl/6J and C3H/HeN adult offspring exhibited higher plasma corticosterone levels during the active period (dark) in comparison with the inactive period (light) (Period effect, F_(1,34)_ = 24.17, p < 0.0001 and F_(1,36)_ = 71.01, p < 0.0001, in C57Bl/6J and C3H/HeN respectively). In C3H/HeN males, there was a significant period x MS effect (Figure 2B, F_(1,36)_ = 6.31, p = 0.0234), and the post-hoc analysis revealed that plasma corticosterone levels were significantly higher in MS C3H/HeN during the dark phase (p = 0.0086). In C57Bl/6J mice there was an overall increase of plasma corticosterone in MS animals (Figure 2A, MS effect, F_(1,34)_ = 4.102, p = 0.0508), but this effects was mainly observed during the dark period (p = 0.0213). A similar increase of plasma corticosterone during the dark phase was observed in female offspring exposed to MS (Figure S1E and S1F). This effect was reported in both strains (C57Bl/6J, Control versus MS, p = 0.0032; C3H/HeN strain, control versus MS, p = 0.0331). Regarding other metabolic hormone levels, MS significantly increased plasma resistin levels in the C57Bl/6J strain (Figure 2C, t_(23)_ = 2.640, p = 0.0146), but no difference was observed in C3H/HeN strain (Figure 2D). Finally, in both strains, MS had no effect on insulin and leptin levels (Figure 2E-H).

**Figure 2.**
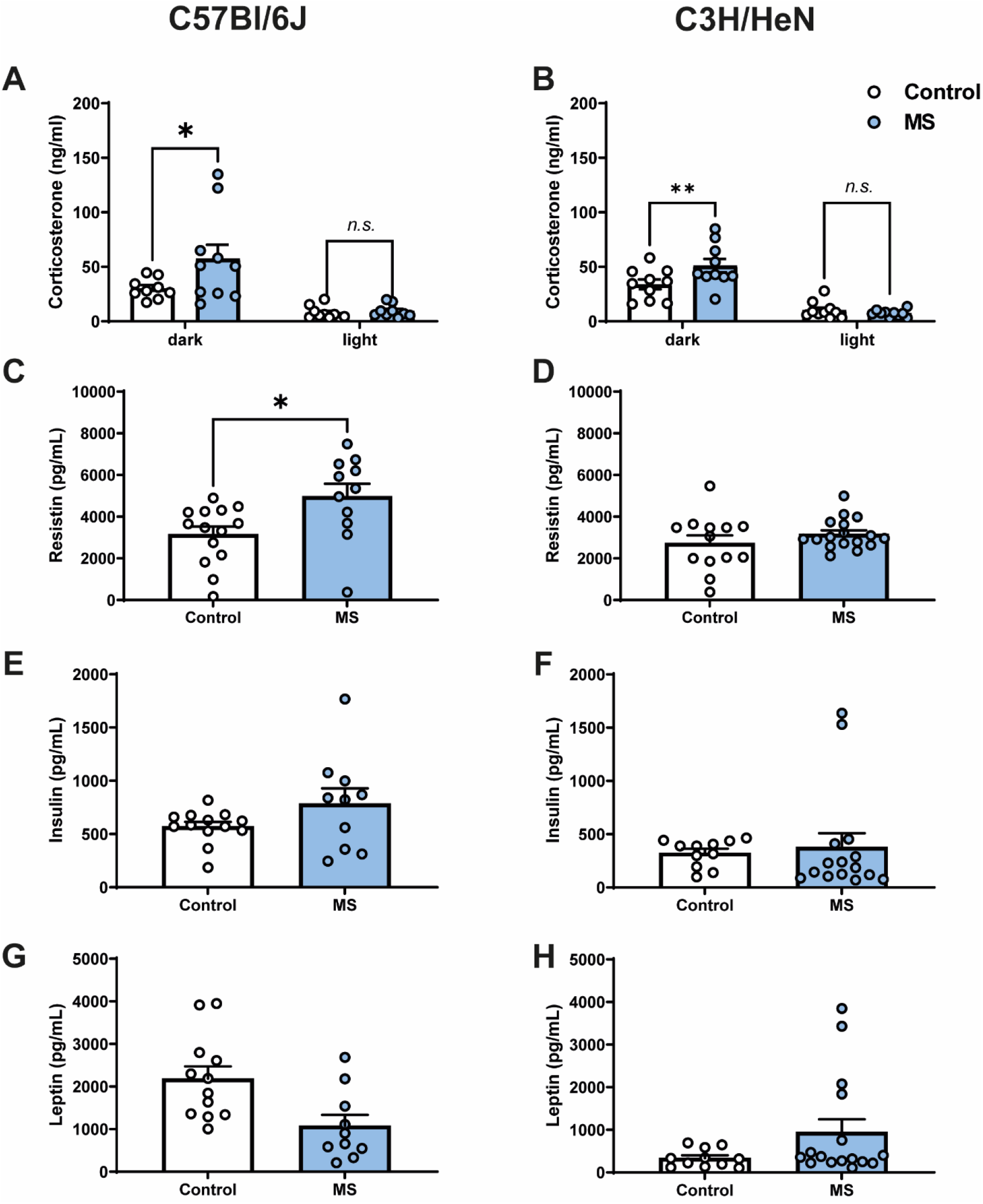
Maternal separation increases plasma corticosterone levels in C57Bl/6J and C3H/HeN male offspring. Plasma corticosterone levels in Control and MS male offspring on a reverse light cycle in both light and dark periods in (A) C57Bl/6J (Control n = 9, MS n = 10) and (B) C3H/HeN (Control n = 10, MS n = 10). Plasma levels in control and MS male offspring after a 12-hr fast of insulin in (C) C57Bl/6J (Control n = 13, MS n = 10) and (D) C3H/HeN (Control n = 11, MS n = 15), resistin in (E) C57Bl/6J (Control n = 14, MS n = 11) and (F) C3H/HeN (Control n = 13, MS n = 17), and leptin in (G) C57Bl/6J (Control n = 12, MS n = 11) and (H) C3H/HeN. Data are presented as means ± SEM, *p < 0.05, ***p < 0.01.

### 3.3 Chronic maternal separation exacerbates the motivation for palatable food of C3H/HeN mice fed *ad libitum*

We used operant conditioning paradigm to examine the effect of MS on the motivation for palatable food. During the initial training on a fixed-ratio 1 (FR-1) schedule (with mild food deprivation), both strains similarly increased their number of presses on the active lever across sessions indicating that MS did not affect instrumental learning (sessions effect, C57Bl/6J strain: F_(4,110)_ = 16.15, p < 0.0001 Figure 3A; C3H/HeN: F_(4,110)_ = 39.41, p < 0.0001, Figure 3B). The lever presses on the inactive lever were very low and similar between groups. When mice were fed *ad libitum* (5th session under FR1 schedule), the number of presses on the active lever was decreased, but control and MS groups showed similar behaviour. These results indicated that MS did not alter the learning in this task and that both strains maintained a motivation for palatable food without any food restriction.

**Figure 3.**
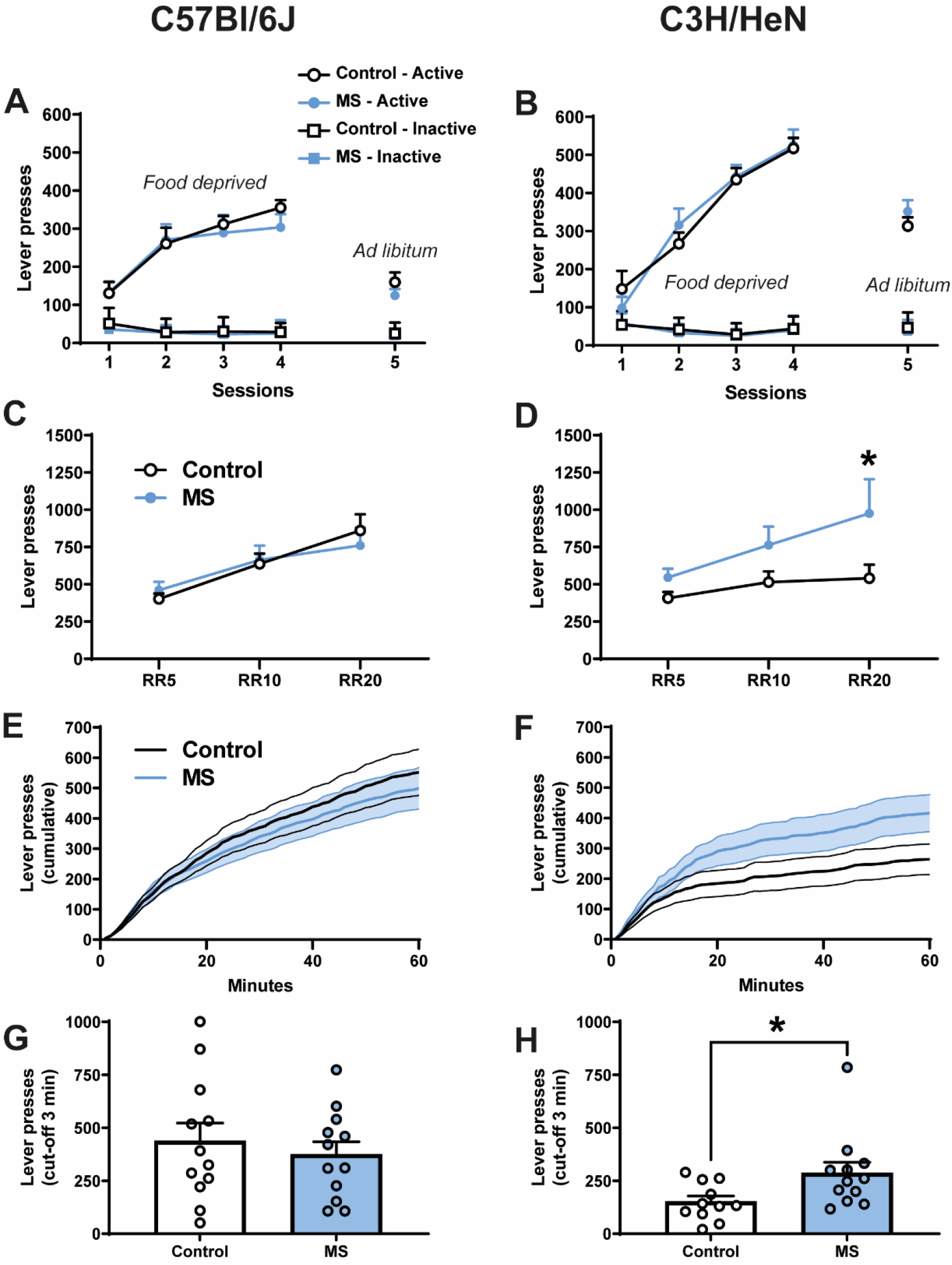
Motivation for palatable food is exacerbated in male C3H/HeN mice submitted to maternal separation. Fixed-ratio 1 schedule with the number of lever presses over the first 4 food deprived sessions and the last 5^th^ *ad libitum* session comparing control and MS mice in both (A) C57Bl/6J and (B) C3H/HeN. r ratio schedule (probability of 1/5, RR5; 1/10, RR10; 1/20, RR20) with number of lever presses in Control and MS mice in both (C) C57Bl/6J and (D) C3H/HeN. Progressive-ratio ×2 schedule with cumulative number of lever presses over a 60-min session in Control and MS mice in both (E) C57Bl/6J and (F) C3H/HeN, and the number of lever presses before a 3-min cut-off in both (G) C57Bl/6J and (H) C3H/HeN. Data are presented as means ± SEM, *p < 0.05, **p < 0.01. n = 11-12 mice/group.

A random ratio (RR) schedule with a 1/5, 1/10 and 1/20 probability (RR5, RR10 and RR20) to obtain one food reward was next performed in *ad libitum* fed animals (Figure 3C and D, for C57Bl/6J and C3H/HeN, respectively). In both strains, the number of active lever presses increased according to the probability to obtain the reward (RR effect, F_(2,22)_ = 47.16, p < 0.0001 for C57Bl/6J and F_(2,22)_ = 9.606, p = 0.0020 for C3H/HeN). However, while MS and control animals did not differ in the C57Bl/6J strain (MS effect, F_(1,22)_ = 0.0021, n.s.), MS C3H/HeN mice showed a tendency for an overall increase of their active lever presses (MS effect, F_(1, 22)_ = 3,5042, p = 0,07457). The MS effect was significant in C3H/HeN mice for the RR20 schedule when the effort required to obtain the food reward was the highest (Figure 3D, p = 0.0386). A similar profile was reported in females with no impact of MS in C57Bl/6 (Figure S1G) and a significant increase of lever presses in C3H/HeN mice exposed to MS (Figure S1H, MS effect, F_(1, 13)_ = 5.818, p = 0.0314).

Under a progressive ratio schedule (PRx2), control and MS C57Bl/6J mice showed similar lever presses profiles (Figures 3E and 3G); in contrast, C3H/HeN exposed to MS had a tendency to increase their overall total amount of lever presses over a 1-h session (t_(22)_ = 1.971, p = 0.0683) and significantly increased their number of lever presses before the cut-off (t_(21)_ = 2.278, p = 0.0333) compared to the control group. In females, whatever the strain, MS had no significant impact (data not shown), indicating that only MS male C3H/HeN showed an exacerbated motivation for palatable food.

In order to determine whether the exacerbated motivation in MS male C3H/HeN was due to an alteration of the representation of outcome value, we studied their performances in a devaluation procedure. In the sensory-specific satiety test, a significant effect of pre-feeding was observed on the number of lever presses in both strains exposed or not to MS (Figure 4A, C57Bl/6J, devaluation effect, F_(1,22)_ = 6.98, p = 0.0148 and Figure 4B, C3H/HeN, F_(1,22)_ = 9.23, p = 0.0060). Finally, in the concurrent choice test (conducted in fasted animals), the presence of an alternative reward less palatable (chow), but freely available in the operant chamber, reduced the number of lever presses for sweetened milk similarly between control and MS groups of both strain (Figure 4C, C57Bl/6J, (F_(1,22)_ = 131.9, p < 0.0001; Figure 4D, C3H/HeN, F_(1,22)_ = 80.75, p < 0.0001). The total amount of chow consumed during the test was similar between groups in both strains (Figure 4EF).

**Figure 4.**
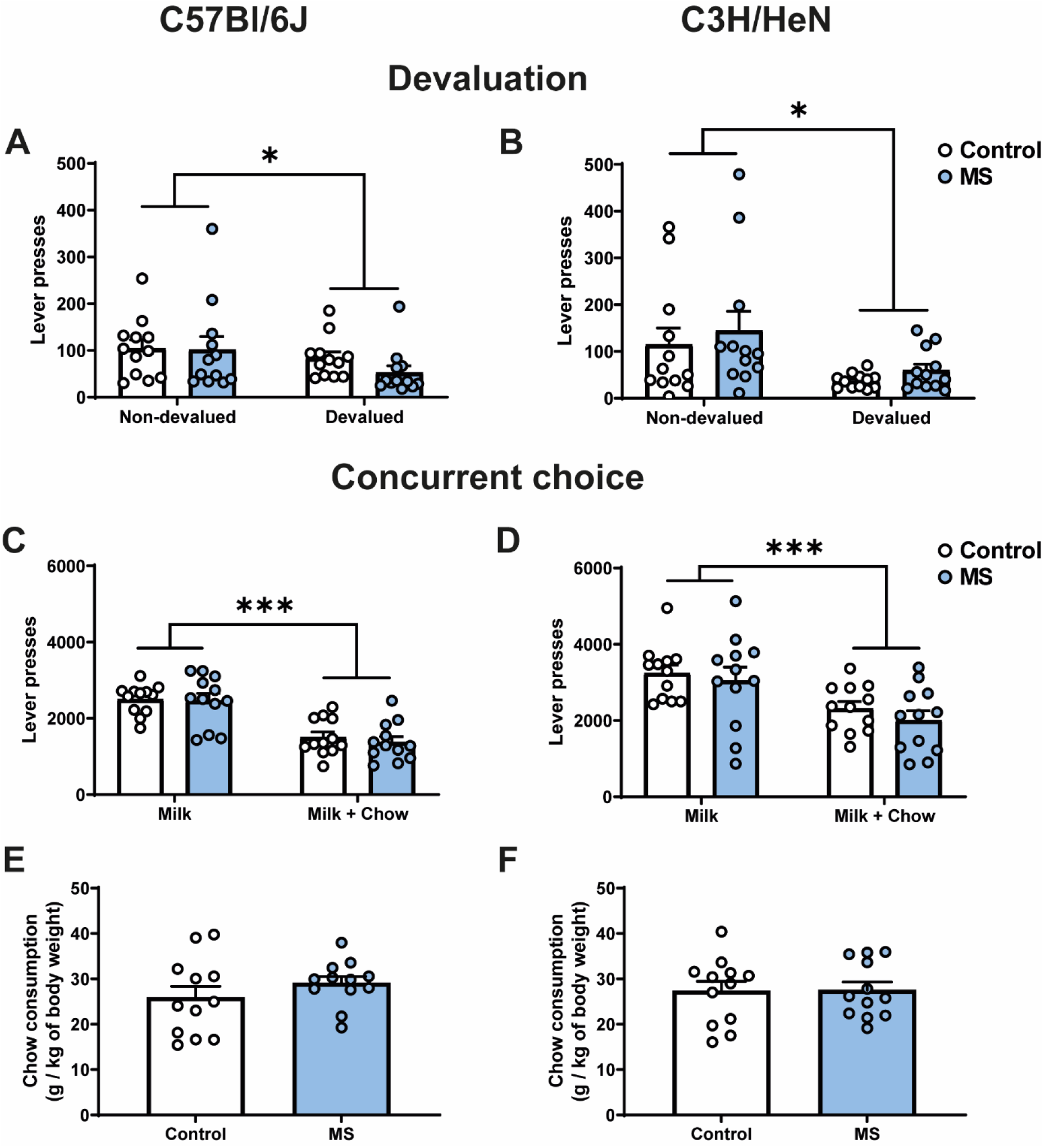
Maternal separation does not affect representation of outcome value and rate of lever presses in the concurrent choice procedure. Lever presses associated with Milk or Intralipid after a RR20 schedule presses comparing Control and MS mice in both strains ((A1) C57Bl/6J and (A2) C3H/HeN, respectively). Presses on lever under a concurrent choice task between control and MS mice in (B1) C57BL/6J and (B2) C3H/HeN. Chow consumption comparing Control and MS mice in (B3) C57BL/6J and (B4) C3H/HeN. Data are presented as means ± SEM. n = 12 mice/group.

### 3.4 Maternal separation modifies the transcriptional profile in Nucleus Accumbens of C3H/HeN

To investigate the molecular brain signature associated with exacerbated motivation in MS animals, after the operant task, we performed transcriptomic analysis in male C3H/HeN offspring. We hybridized RNA from whole-tissue micropunches of NAc, mPFC, and hypothalamus on Agilent microarrays. Descriptive analysis *via* hierarchical clustering showed a clear separation for NAc only (Figure 5A, Figure S2A and D). PLS-DA analysis was able to build a model in each tissue but only the model in NAc was validated (Figure 5B, *p*R2Y = 0.014, *p*Q2 = 0.02 for NAc, Figure S2B *p*R2Y = 1, *p*Q2 = 0.84 for mPCF and Figure S2E *p*R2Y = 0.78, *p*Q2 = 0.94 for hypothalamus). No differentially expressed genes (DEGs) were found in mPFC and hypothalamus (Figure S2C and F), while 375 DEGs were found in NAc: 306 downregulated and 69 upregulated (Figure 5C). Therefore, the transcriptional effect is restricted to the NAc tissue. A bioinformatic analysis of functions and networks was carried on the 375 DEGs using the IPA software and the ConsensusPathDB-mouse (CPDB) website. This detected an overrepresentation of 8 canonical pathways with BH *p-*values < 0.05 and 18 KEGG pathways with *q-*values < 0.05, respectively (Figure 5D). Among these pathways, 2 are common to the 2 analyses: GABA receptor signalling/GABAergic synapse and Glutamate receptor signalling/Glutamatergic/synapse. Pathways analysis also pointed out stress related pathways (a-adrenergic signalling and CRH signalling) and addiction pathways (nicotine, morphine cocaine). The IPA analysis also predicted upstream regulators, which may be causing the observed gene expression changes. Interestingly, among the 3 predicted upstream regulators, the L-Dopa had the lowest *p*-value and the highest *z*-score, with a predicted activation whereas the uncoupling protein 1 *Ucp1* and the histone deacetylases *Hdac* were predicted as inhibited (Figure 5E). Finally, using TLDA assays, we performed a RT-qPCR analysis on 84 genes of NAc samples from males and females of both strains. Microarray results obtained in C3H/HeN male mice were validated by TLDA (Figure S2G, Spearman correlation *rho* = 0.754; *p* < 2.2e^-16^, supplementary file F1). A clustering analysis of these TLDA results showed clearly the differential expression of the 84 genes between control and MS C3H/HeN males. In contrast, we did not observe any differential expression for the 84 genes identified in C3H/HeN mice when we compared control and MS C3H/HeN females or control and MS C57Bl/6J regardless ofthe sex (Figure S2H, supplementary file F1).

**Figure 5.**
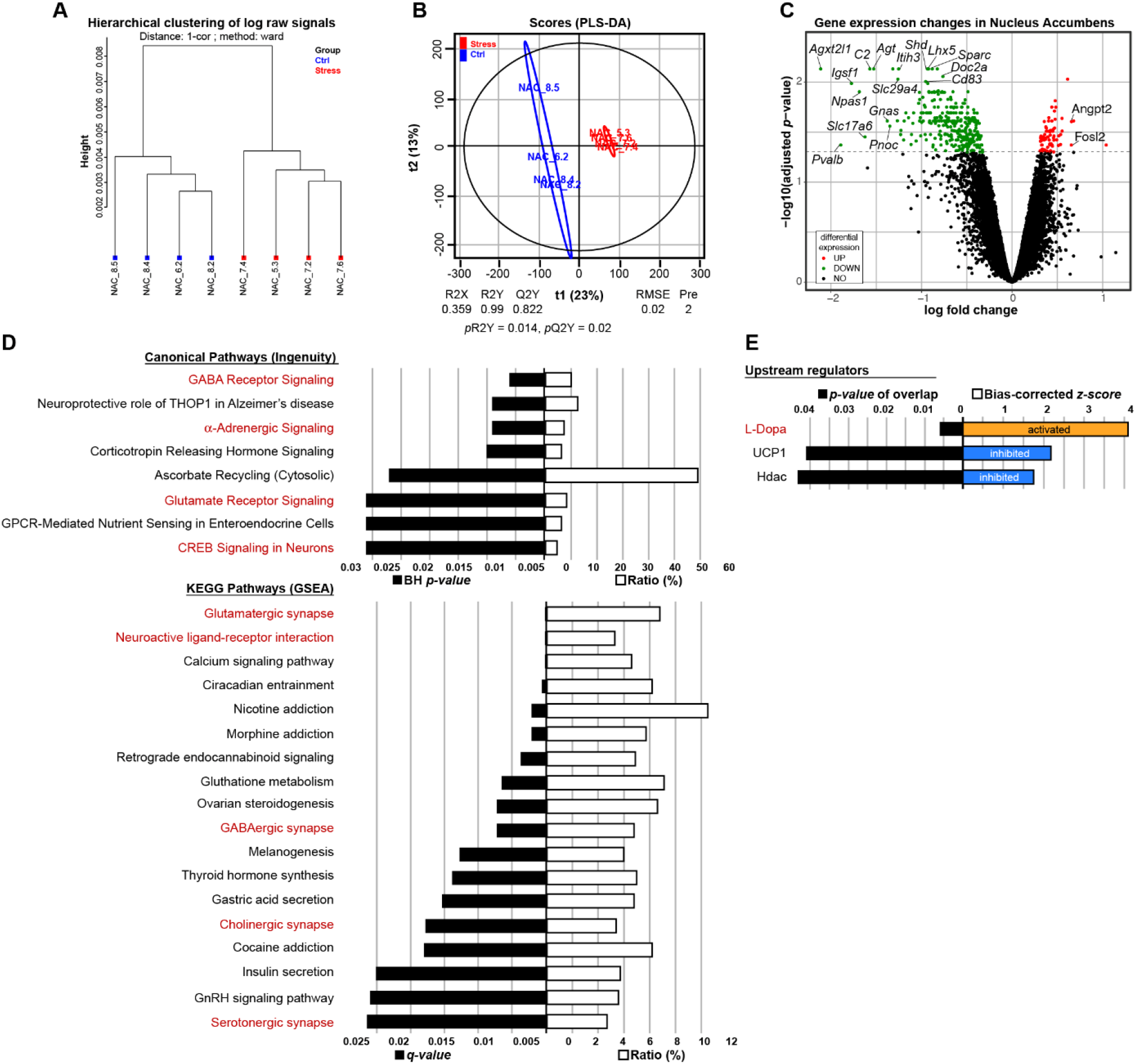
Maternal separation drastically modifies the transcriptional profile in Nucleus Accumbens (NAc) of male C3H/HeN mice. Clustering by Euclidean distance of the raw transcriptomic datasets of NAc from Control (blue) or MS (red) male adult offspring **(A)**. Score plots from the PLS-DA classification into Control (blue) and MS (red) groups. A model is considered robust when the response variance explained (R2Y = 0.99) is higher than the predictive performance of the model (Q2Y = 0.822). A model with a Q2Y > 0.5 is considered to have a good predictive performance **(B)**. Volcano plot depicting significantly differentially expressed genes in the NAc of C3H/HeN mice between the MS and Control conditions **(C)**. Top Canonical Pathways and KEGG pathways **(D)** and top upstream regulators **(E)** involved. In red are depicted the pathways or regulators associated with NAc neuronal function. n = 4 mice/group.

## 4 Discussion

MS models in rodent have been widely used to examine long lasting effects of early-life stress on emotional function. Here we studied the impact of early-life stress on the motivation for a palatable nutritional reward in two mouse strains. Our data showed that MS associated with unpredictable chronic mild stress in lactating dams exacerbated motivation for a palatable food reward (sweetened milk) in both males and females C3H/HeN mice. Interestingly, no effect of MS on C57Bl/6J mice strain’s motivation was observed. The transcriptomic analysis revealed that exacerbated motivation in MS C3H/HeN male mice was associated with marked changes in gene expression in the NAc, whereas no significant changes were reported in the PFC or hypothalamus. Furthermore, MS C57BL/6J mice which showed the same motivation for palatable food than control mice also exhibited a similar genes expression than control animals.

A primary and key result of our study is the overall difference in behavioural and brain gene expression outcomes observed between C3H/HeN and C57Bl/6J strains in response to early life stress. Despite a significant effect on HPA axis in both strains, body weight and food motivation were specifically altered in C3H/HeN but not in C57Bl/6J suggesting that C3H/HeN strain is more susceptible to MS for these studied parameters. This comforts previous findings showing a resilience of the C57Bl/6 strain to the MS procedure effects (Millstein and Holmes, 2007; Tractenberg et al., 2016). Indeed, despite the fact that MS procedure has been fairly well-established in rats (O’Mahony et al., 2009; Rincel and Darnaudéry, 2020), results in mice especially in the C57Bl/6 strain are very heterogenous. Previous work using an early life stress paradigm close to our study (3 hrs MS from PND1 to PND14, coupled to an unpredictable restraint stress or forced swim stress in dams during separation) showed increased depressive-like behaviours, reduced anxiety, but no impact on serum insulin levels in C57Bl/6 offspring exposed to early stress (Franklin et al., 2010; Gapp et al., 2014; van Steenwyk et al., 2018). Regarding HPA axis function, we demonstrated that MS increases plasma corticosterone at the beginning of the dark phase in both strains, this being coherent with previous results in the literature (Tractenberg et al., 2016; Zajdel et al., 2019). This result is important since it demonstrates that despite differential strains’ sensitivity to MS in food motivation, both strains are affected by MS. Most of previous works conducted in C57Bl/6 exposed to MS report long-term effects on behaviour, only when animals were re-exposed to an additional stressor such as early weaning at PND17 (Tchenio et al., 2017), or chronic social defeat or UCMS at adulthood (Peña et al., 2017, 2019). Without additional stress, emotional behaviours in MS C57Bl/6 mice generally did not significantly differ from controls (Millstein and Holmes, 2007; Tractenberg et al., 2016; Peña et al., 2017, 2019; Baugher and Sachs, 2022; Shin et al., 2023). On the other hand, C3H/HeN strain used in the present study has been previously reported to induce gut dysfunction after MS (Riba et al., 2018; Ilchmann-Diounou et al., 2019; Rincel et al., 2019) and emotional impairments in a multi-hit model combining MS, maternal UCMS and prenatal infection (Rincel et al., 2019). The different susceptibility to MS procedure between the C57Bl/6J and C3H/HeN strains could be due to maternal care differences. Accordingly, C3H/HeN dams display more pups licking and nursing than other strains (van der Veen et al., 2008). This robust maternal behaviour may be more affected by the disruption of the nest and pups’ separation associated to MS procedure. Additionally, C3H/HeN strain are more anxious and show a higher sensitivity to stress compared to other mice strains; whereas C57Bl/6J strain has been recurrently described in the literature as a strain particularly resilient to stress (Moloney et al., 2015). Taken together, this suggests that MS effects may vary according to both the strain and the dimension considered (endocrine responses, metabolism, behaviours: emotion, cognition, or motivation).

Although emotional behaviours have been extensively studied in MS literature, motivation for food reward has been less explored. Here, we present data supporting an impact of early-life stress on palatable food motivation. Previous works in rodents exposed to early-life stress, demonstrate that MS exacerbates drugs of abuse motivation and ethanol intake (Matthews et al., 1999; Walters and Kosten, 2019). Since drug and palatable food rewards partially share common neurobiological substrates, we hypothesized that MS will affect food motivation and impact mesolimbic circuit. Previous findings in rats showed that early-life stress exacerbates motivation for palatable food as indicated by their higher breakpoint in operant task and their lower latency to reach chocolate pellets in a runway task (Romaní-Pérez et al., 2017; Levis et al., 2021). Here, we report for the first time an increased motivation for a palatable food reward in an operant paradigm in mice after early-life stress exposure. Interestingly, we also report a marked strain effect, as this effect was specifically observed in the MS C3H/HeN mice, but not in MS C57Bl/6J mice. This effect is observed in both male and female C3H/HeN offspring exposed to MS especially in RR20 and PR schedules when the effort to obtain the food reward is high. It is important to note that here, exacerbated motivation for palatable food is reported in mice non-submitted to food restriction during the instrumental task. These results suggest that independently of a nutritional status, stressed animals may be more prone to seek and consume calorie-dense food. In that way, MS rats ate more palatable food when they have a free access in their home cage (Romaní-Pérez et al., 2017; de Lima et al., 2020; Levis et al., 2021). Our results are in accordance and extend recent finding showing that a single long lasting (23 h) separation at PND3 produces enhanced binge-eating behaviour after repetitive cycles of reexposure to a high-fat diet in adulthood (Shin et al., 2023) and clinical literature showing that early-life adversity is associated with higher risk to develop food addiction behaviours and obesity in adulthood (Rorty et al., 1994; Mason et al., 2013; Danese and Tan, 2014).

An important finding in the present work is that MS C3H/HeN males exhibiting exacerbated motivation for palatable food had marked changes of brain gene expression (adj p-values: 375 genes differentially expressed) specifically in the NAc, a key region for the regulation of motivation and reward processing. Notably, we validated a large amount of gene differentially expressed in MS C3H/HeN males using TLDA. Interestingly none of the genes significantly affected by early-life stress in C3H/HeN mice were changed in C57Bl/6J MS mice. However, given that C57Bl/6J results were not obtained using microarray, we cannot exclude that changes also occur in this group but affecting other genes. Previous transcriptomic studies demonstrated a significant impact of early-life stress using RNAseq analysis in C57Bl/6 strain, although authors used non-adjusted p-values (Peña et al., 2017, 2019). Using adjusted p-values, we did not detect a significant effect of MS on gene expression in the PFC and hypothalamus, indicating that NAc is a brain area particularly affected by MS in C3H/HeN. The lack of transcriptional change within the hypothalamus is quite surprising considering the importance of this brain area in the effects of stress and in the control of food intake. Again, it is important to note that we used here the Benjamini-Hochberg adjusted p-values method which is highly conservative and may lead to under detection of change in gene expression between groups. Further studies should be conducted to examine the impact of MS on specific nuclei of the hypothalamus such as lateral hypothalamus. Pathways analysis (using IPA and CPDB) on NAc genes revealed that the excitatory (Glutamate receptor signalling) and inhibitory (GABA_A_ receptor signalling) pathways are both affected by early-life stress. Gabaergic and glutamatergic systems play a major role in neurodevelopment and disruption of these systems have been involved in numerous neurodevelopmental disorders including autism, schizophrenia or ADHD (Zhang et al., 2020). Furthermore, a large body of evidence has linked various perinatal stress paradigms with altered excitatory/inhibitory balance (Holland et al., 2014; Karst et al., 2020; Ohta et al., 2020; Crombie et al., 2022). We identified several genes such as *Agt, Igsf1, Gnas, Pnoc, Npas1, Pvalb* or *Fosl2* affected by early life stress in C3H male mice which have been previously reported to be modified in the NAc or in the VTA after chronic stress procedures (Peña et al., 2017, 2019). Interestingly, *Gnas* (Guanine Nucleotide-Binding Protein G Subunit Alpha) and *Pnoc* (prepronociceptin) have previously been linked to motivation and reward regulation (Zhu et al., 2016; Parker et al., 2019; Rodriguez-Romaguera et al., 2020; Liu et al., 2022). *Angpt2* (angiopoietin-2) plays a role in angiogenesis and its expression is induced by inflammatory markers (Akwii et al., 2019). *Angpt2* has not been linked with motivational changes, but recent data suggest that up-regulation of immune factors within the NAc may correlate with addictive phenotype in rats (Calipari et al., 2018) and compulsive sucrose seeking in mice fed with high-fat diet (Décarie-Spain et al., 2018). Importantly, major upstream regulators identified by IPA are *L-Dopa, Ucp1* and *Hdac* suggesting that these factors may contribute to the effects of early stress. L-Dopa is a precursor for dopamine. Interestingly, NAc dopamine plays a critical role in palatable food-seeking behaviour for reinforcement schedules that have high-work requirements such as RR20 or PR (Salamone et al., 2007). HDAC is an important epigenetic regulator and its inhibition in the NAc promotes drug self-administration (Wang et al., 2010). Finally, UCP1 (uncoupling protein 1, a proton carrier protein generating heat via non-shivering thermogenesis) has been recently identified in the brain, though its role still remains unclear (Claflin et al., 2022).

In conclusion, our work reveals that early-life stress increases motivation for palatable food in *ad libitum* fed adult C3H/HeN mice. This effect is associated with marked changes in gene expression within the NAc. Importantly, exacerbated palatable food motivation after MS was reported in both male and female C3H /HeN mice, but this effect is strain dependent suggesting a relative resilience of C57Bl/6. Overall, our study confirms that early-life adversity has enduring effects on reward circuits and highlights the importance to further explore the impact of early-life stress on motivational processes.

## Acknowledgements

We thank Claire Naylies and Yannick Lippi for their contribution to microarray fingerprints acquisition and microarray data analysis carried out at GeT Genopole Toulouse Midi-Pyrénées facility (https://doi.org/10.15454/1.5572370921303193E12). This work was supported by Agence Nationale de la Recherche (ANR-12-DSSA-0004 Incorporation Biologique et Inégalités Sociales de Santé) ; Fondation recherche médicale (ENV-Environnement – Santé ; ENV202003011543) ; Université de Bordeaux and INRAE.

---

**Figure S1.**
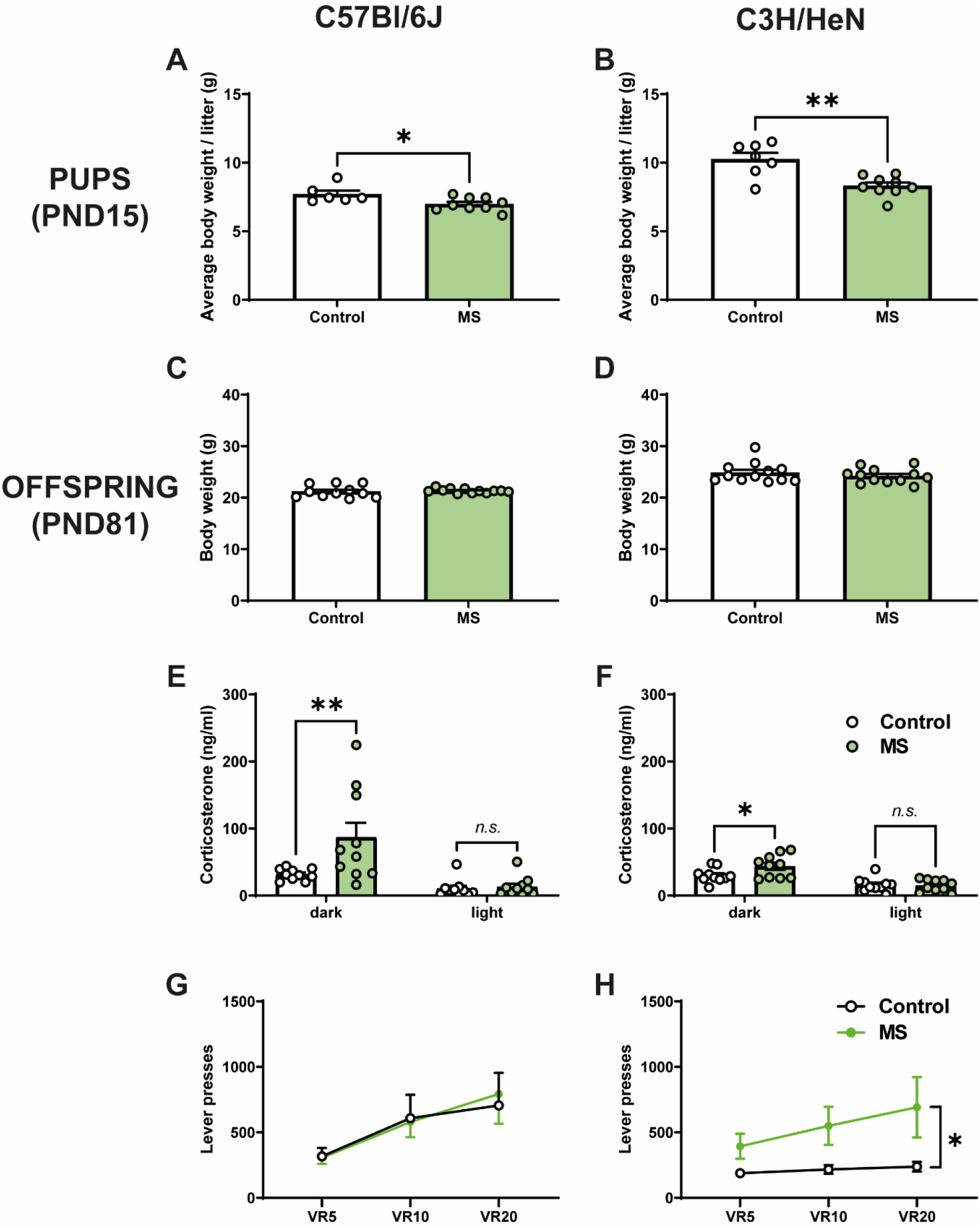
Maternal separation impairs pup’s body weight, corticosterone level and motivation for palatable food in female C3H/HeN offspring. Body weight in Control and MS animals in female (A) C57Bl/6J pups at PND15, (B) C3H/HeN pups at PND15, (C) C57Bl/6J offspring at PND81, and (D) C3H/HeN offspring at PND81. Plasma corticosterone levels in Control and MS female offspring on a reverse light cycle in both light and dark periods in (E) C57Bl/6J and (F) C3H/HeN. Random ratio schedule (probability of 1/5, RR5; 1/10, RR10; 1/20, RR20) with number of lever presses in control and MS female offspring in (G) C57Bl/6J and (H) C3H/HeN. Data are presented as means ± SEM, *p < 0.05, **p < 0.01. n = 7-10 mice/group.

**Figure S2.**
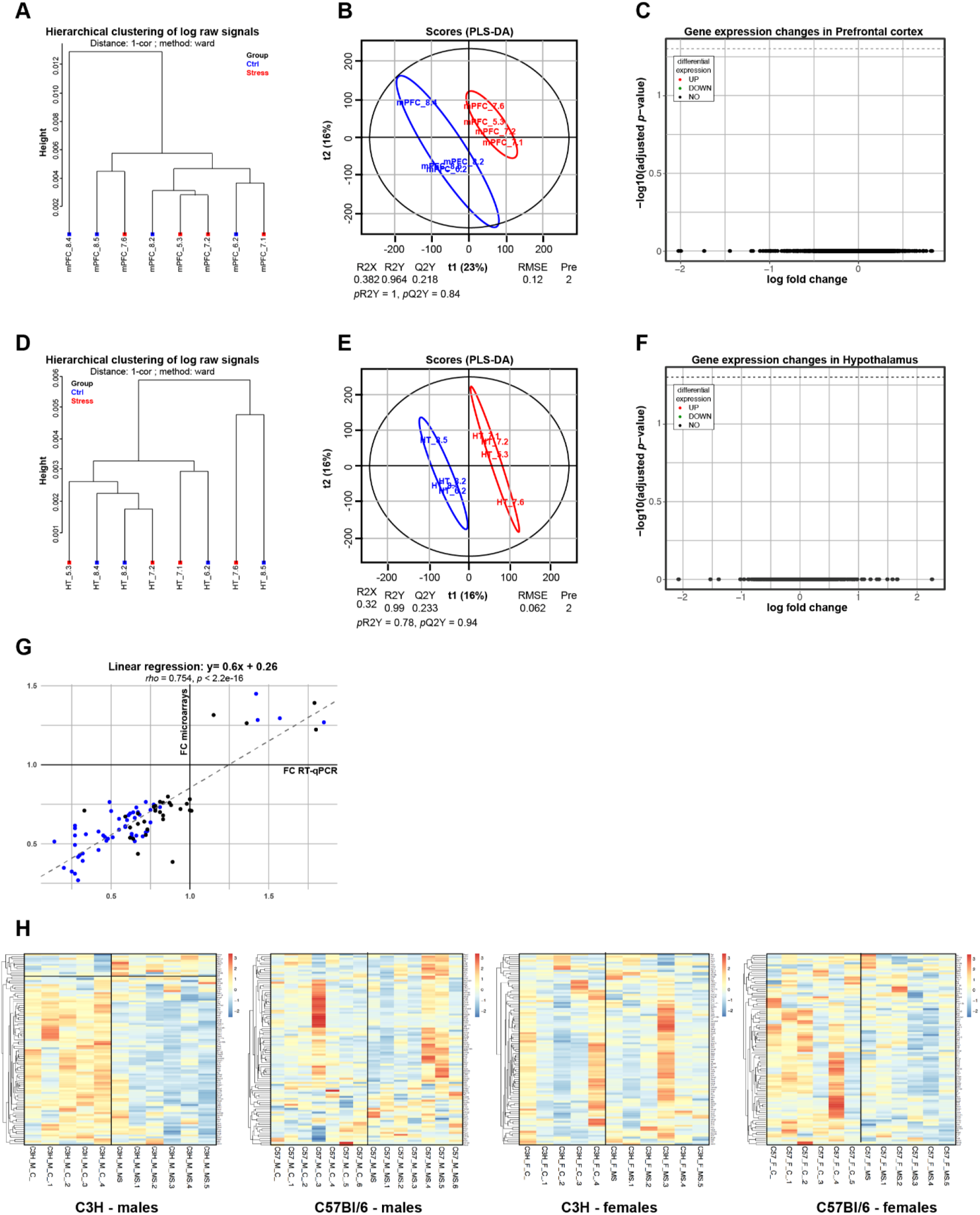
Additional information concerning the transcriptomic analysis. Clustering by Euclidean distance of the raw transcriptomic datasets from Control (blue) or MS (red) male adult offspring of mPFC **(A)** and HT **(D)**. Score plots from the PLS-DA classification into Control (blue) and MS (red) groups. A model is considered robust when the response variance explained (R2Y) is higher than the predictive performance of the model (Q2Y). A model with a Q2Y > 0.5 is considered to have a good predictive performance, which is not the case neither for mPFC **(B)** not HT **(E)**. Volcano plot depicting no significantly differentially expressed genes were found between the MS and Control conditions neither for mPFC **(C)** not HT **(F)**. n = 4 mice/group. A plot of the fold change (FC) calculated for RT-qPCR and microarray values between male MS and Control NAc for 84 genes. The RT-qPCR data correlated with the microarray data for most of the 84 FC, with a linear equation: y = 0,6x + 0.26 and a high correlation coefficient (rho = 0.754). Spearman’s test indicated that this correlation was highly significant (p < 2.2e10-16). More than 60% of the genes tested (51 out of the 84) were significantly differentially expressed (adj p < 0.05, blue dots) n = 4 mice/group **(G)**. Expression clustering for the 87 gene studied in the TLDA for C3H/Hen males (n=5 control, 6 MS), C57Bl/6J males (n = 7 control, 7 MS), C3H/HeN females (n = 5 control, 6 MS) and C57Bl/6J females (n = 6 control, 6 MS).

## Notes

### Competing Interest Statement

The authors have declared no competing interest.

